# Conformational entropy and specific affinity of M-cholinolytics and ligands of µ-opioid and D_2_-dopamine receptors

**DOI:** 10.1101/2020.04.06.026302

**Authors:** Mikhail B. Darkhovskii, Felix S. Dukhovich

## Abstract

The computation model for evaluation of conformational entropy changes upon binding ligands to receptors is described. Then, changes of conformational entropy component and of binding free energy are compared. Interest to conformational entropy arises from developing new drugs as it might be changed purposefully. It is shown that conformational entropy may be used for prediction of affinity to a certain receptor. Examples of directed affinity change under the modification of substances’ conformational flexibility are given. The specific role of the conformational entropy in the receptor’s protection from the irreversible inactivation is identified.

## Introduction

There are known problems in computation of conformational entropy. First, conformational entropy contribution is often implicit, as modification of compounds structure is accompanied by changing many enthalpic and entropic factors, besides the conformational entropy changes. Second, only the total entropy change of the ligand-receptor complex formation can be determined in experiments. The entropy change during ligand-receptor complex formation is a sum of the following terms [1]:

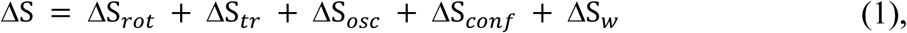

where the first four terms are respectively from rotational, translational, vibrational, and conformational entropy changes of the ligand and the receptor. Fifth term is the entropy change during water displacement (the total or partial removal of hydrate shells) from the receptor’s binding site.

## Substances Selection

The components of the entropy change in Eq.(1) can be estimated by semi quantitative and empirical methods. However, we are interested in such a model that can extract ‘pure’ conformational contribution to the entropy change. That is, the change in the structure of substances should minimize change in the enthalpy and other than conformational components of the entropy. As such substances, it is appropriate to choose the amino esters of substituted glycolic and acetic acids (Table 1).

**Table 1.**
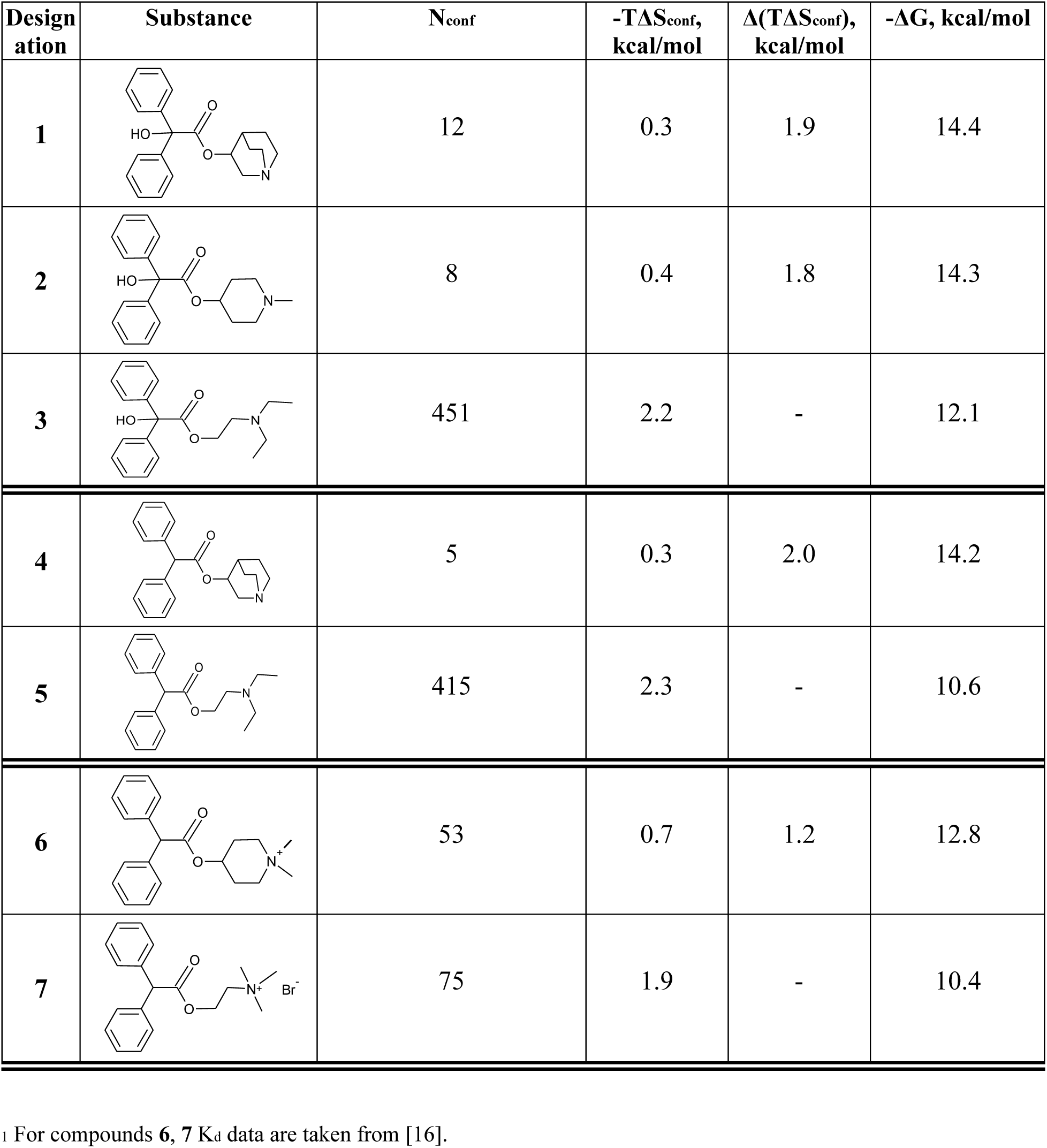
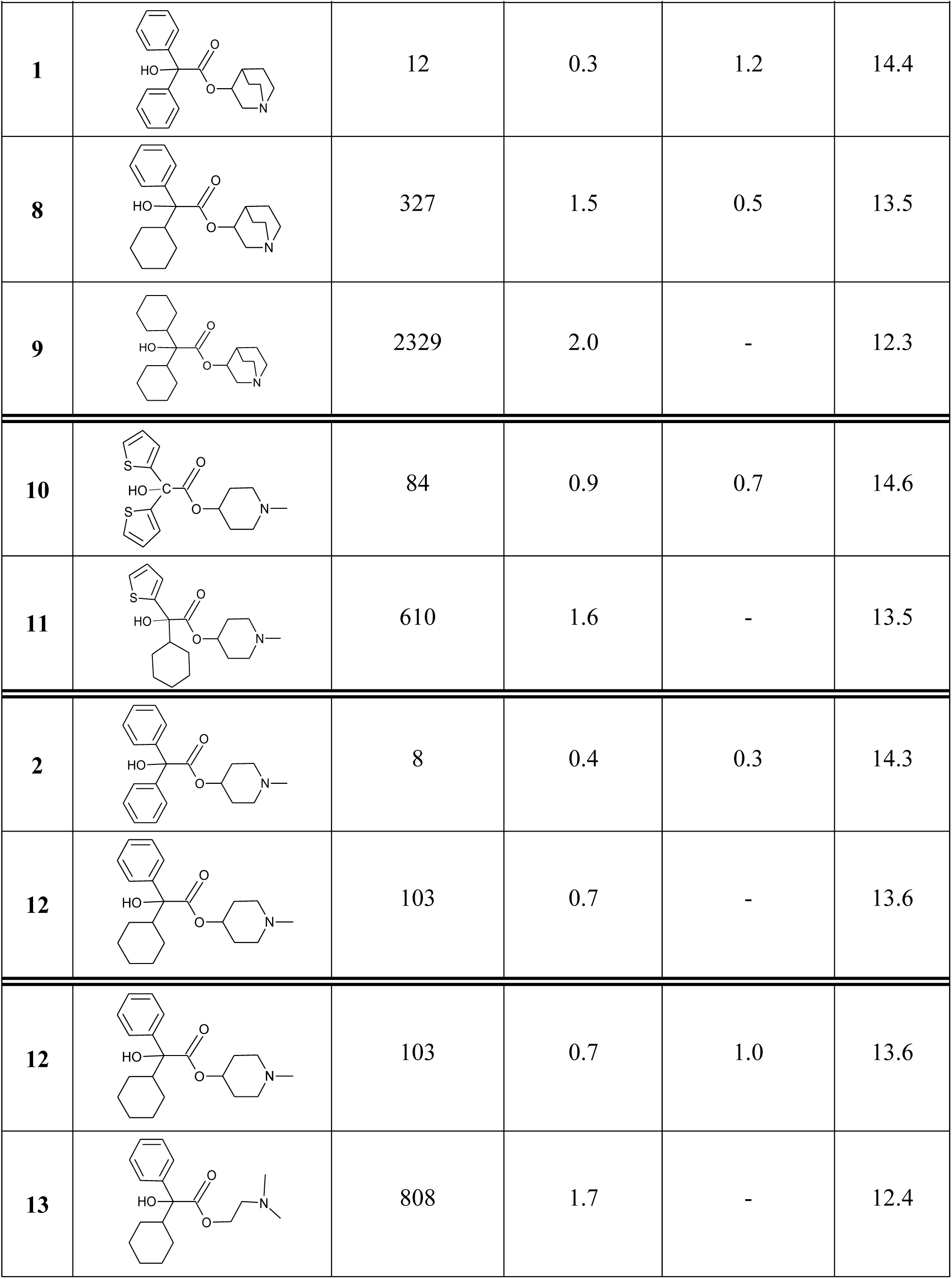
Number of conformations and conformational entropy changes on structural modification of substances^1^.

| Designation | Substance | N<sub>conf</sub> | -TΔS<sub>conf</sub>, kcal/mol | Δ(TΔS<sub>conf</sub>), kcal/mol | -ΔG, kcal/mol |
| --- | --- | --- | --- | --- | --- |
| **1** |  | 12 | 0.3 | 1.9 | 14.4 |
| **2** |  | 8 | 0.4 | 1.8 | 14.3 |
| **3** |  | 451 | 2.2 | - | 12.1 |
| **4** |  | 5 | 0.3 | 2.0 | 14.2 |
| **5** |  | 415 | 2.3 | - | 10.6 |
| **6** |  | 53 | 0.7 | 1.2 | 12.8 |
| **7** |  | 75 | 1.9 | - | 10.4 |

Table 1 (continued)
| Designation | Substance | N<sub>conf</sub> | -TΔS<sub>conf</sub>, kcal/mol | Δ(TΔS<sub>conf</sub>), kcal/mol | -ΔG, kcal/mol |
| --- | --- | --- | --- | --- | --- |
| 1 |  | 12 | 0.3 | 1.2 | 14.4 |
| 8 |  | 327 | 1.5 | 0.5 | 13.5 |
| 9 |  | 2329 | 2.0 | - | 12.3 |
| 10 |  | 84 | 0.9 | 0.7 | 14.6 |
| 11 |  | 610 | 1.6 | - | 13.5 |
| 2 |  | 8 | 0.4 | 0.3 | 14.3 |
| 12 |  | 103 | 0.7 | - | 13.6 |
| 12 |  | 103 | 0.7 | 1.0 | 13.6 |
| 13 |  | 808 | 1.7 | - | 12.4 |

These mAChR antagonists have the similar molecular weight and size, charge distribution, basicity of the amino group, and the same pharmacophore [2]. In addition, anchoring substituents in the acidic part of molecules have similar partial free energy contributions (ΔΔG) to the mAChR binding free energy: phenyl, thienyl, and cyclohexyl give 2.80, 2.95, and 2.40 kcal/mol, respectively [2, 3]. These substances possess high selectivity to muscarinic acetylcholine receptors, not to their specific subtypes [4].

## Methods

### Determination of K_d_ (ΔG) values upon M-cholinolytic – mAChR complex formation

All experimental ΔG values given in Table 1 are determined pharmacologically by suppressing contraction of rat small intestine tissue induced by acetylcholine at 37.5 °C (310.5 K). In experiments the equilibrium dissociation constant K_d_ of the ‘substance – receptor’ complex was determined. It is related to Gibbs free energy by:

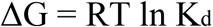

Detailed description of the method can be found in [2, 3, 5]. All investigated substances are synthesized as racemates, so ΔG values of the chiral substances are given in Table 1 for their (R)- or (R,R)-form, for better comparison with non-chiral substances. It is known that affinities of (R)-isomers of the chiral amino esters of substituted glycolic and acetic acids greatly dominate that of (S)-isomers [4]. Due to possible differences in the binding mode, glycolic and acetic acid’ derivatives are considered separately.

### Calculation of the conformational entropy change

To calculate conformational entropy, we need to estimate the number of conformers. TINKER molecular modeling program [6] is used to produce the conformations set. As for all molecules under study, the conformational analysis procedure is based on searching for the potential energy minima in main vibrational modes that correspond to changes in the certain torsion angles. To calculate the potential energy, the MM3 parametrization is used [7]. All the potential energy components used in this parameterization are considered. Calculations were performed for the cationic form in which molecules bind to the receptor.

In order to scan the potential energy surface, we study changes in the torsion angles along the main chain in the ester fragment of glycolates and acetates, substituents at Cα atom of the acidic part, and alkyl groups containing the nitrogen atom. In these settings, we generate conformations at the energy threshold value 10 kcal/mol. The value of ΔS_conf_ is derived based on the assumption that a ligand binds to its receptor at one complementary conformation. Then, the conformational entropy change on the complex formation is expressed through Boltzmann distribution of all rotamers of the ligand [8]:

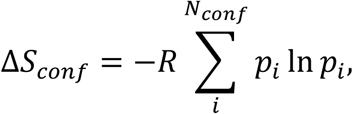

where R is the gas constant, N_conf_ is the number of conformers of an unbound ligand molecule, *i* is the ordinal number of a conformer, and *p*_*i*_ is the ratio of the *i*-th conformer in the set of the conformations, which is expressed at the equilibrium as:

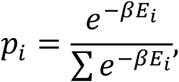

where *E*_*i*_ is the potential energy of the *i*-th conformer, β = 1 / kT (k is the Boltzmann constant). The *p*_*i*_ values are calculated at 310.5 K (37.5 °C), at which the K_d_ values are determined.

Noteworthy, ΔS_conf_ is not proportional to N_conf_, since the *p*_*i*_ values distribution is far from uniform. Conformational changes are not localized in one fragment in which these changes occurred but propagate to the whole molecule. Therefore, modifying one fragment usually cause an increase or decrease in the number of conformations (N_conf_) pertaining to the whole molecule.

## Results and Discussion

Conformational flexibility of the substances is changed in two ways:

- molecular structure framework modification on substitution of more conformationally stable N-methyl-4-piperidyl and 3-quinuclidyl groups for movable N-alkyl group;
- substitution of conformationally rigid phenyl and thienyl for cyclohexyl.

### Conformational entropy and directed affinity change of the substances

Table 1 gives the possibility to compare conformational entropy Δ(TΔS_conf_) and free energy ΔG changes with varied compound’s structural rigidity. The following series of substances were compared: (**1, 2, 3**), (**4, 5**), (**6, 7**), (**1, 8, 9**), (**10, 11**), (**12, 13**), (**2, 12**). An increase in the ligand’s structural rigidity is accompanied by declining of the conformational entropy, and therefore the selected calculation model adequately ‘reacts’ to modifications in the structure of substances. In the first direction, the transition from the relatively labile compound **3** to more rigid **2, 1** reduces the number of conformations N_conf_ from 451 to 8 and 12, and the value of TΔS_conf_ from 2.2 kcal/mol to 0.4 and 0.3 kcal/mol, respectively. Similar structural modifications in acetic acid derivatives (compared ligands **5** and **4**) are accompanied by the reduction of N_conf_ from 415 to 5 and TΔS_conf_ values from 2.3 kcal/mol to 0.3 kcal/mol. Alike pattern of N_conf_ and TΔS_conf_ changes occurs for substances with the quaternary nitrogen atom (**7** and **6**).

The second scenario of the conformational flexibility modification with the substitution of a phenyl group for cyclohexyl is followed by increasing N_conf_ from 12 to 327 and TΔS_conf_ from 0.3 kcal/mol to 1.5 kcal/mol (compounds **1** and **8**). With the replacement of both phenyl rings for cyclohexyl, the N_conf_ sharply rises to 2329 and the TΔS_conf_ to 2.0 kcal/ mol (compounds **1** and **9**). Similar changes occur in (**10, 11**) and (**2, 12**) pair comparison. Thus, conformational entropy might be used to forecast substances affinity to the receptor.

The comparison of the calculated conformational entropy and binding free energy of the pairs of ligands gives interesting result: the following relationship is observed ΔΔ*G*/Δ(*T*Δ*S*_*conf*_) ≈ 2. This suggests the conclusion that conformational entropy losses are distributed approximately equally between the mAChR receptor and the antagonists under study, and that conformational changes in mAChR are localized in the active center of the receptor. However, raising these conclusions to the level of predictability would be premature due to the difference in the methods for determining the change in conformational entropy and free energy. We attribute the result to the successful selection of the calculation model and specific ligands.

It is worth to mention that only two ligands from Table 1, benactyzine (**3**) and adiphenine (**5**), are used as pharmaceuticals [9]. Upper level of drug affinity lies near K_d_ ∼ 0.1 – 0.01 nM. Substances with K_d_ < 0.1 nM cannot be considered as drugs since it follows from the relationship between complexes lifetime and K_d_ value that they almost irreversibly inactivate receptor, so they become poisonous [2, 10].

### Antidepressants and conformational entropy

The conformational entropy dependence on structural rigidity is investigated for other classes of pharmacological agents. In Table 2 the results of N_conf_ and TΔS_conf_ calculations are given for some antidepressants, ligands of D_2_-dopamine receptors.

**Table 2.**
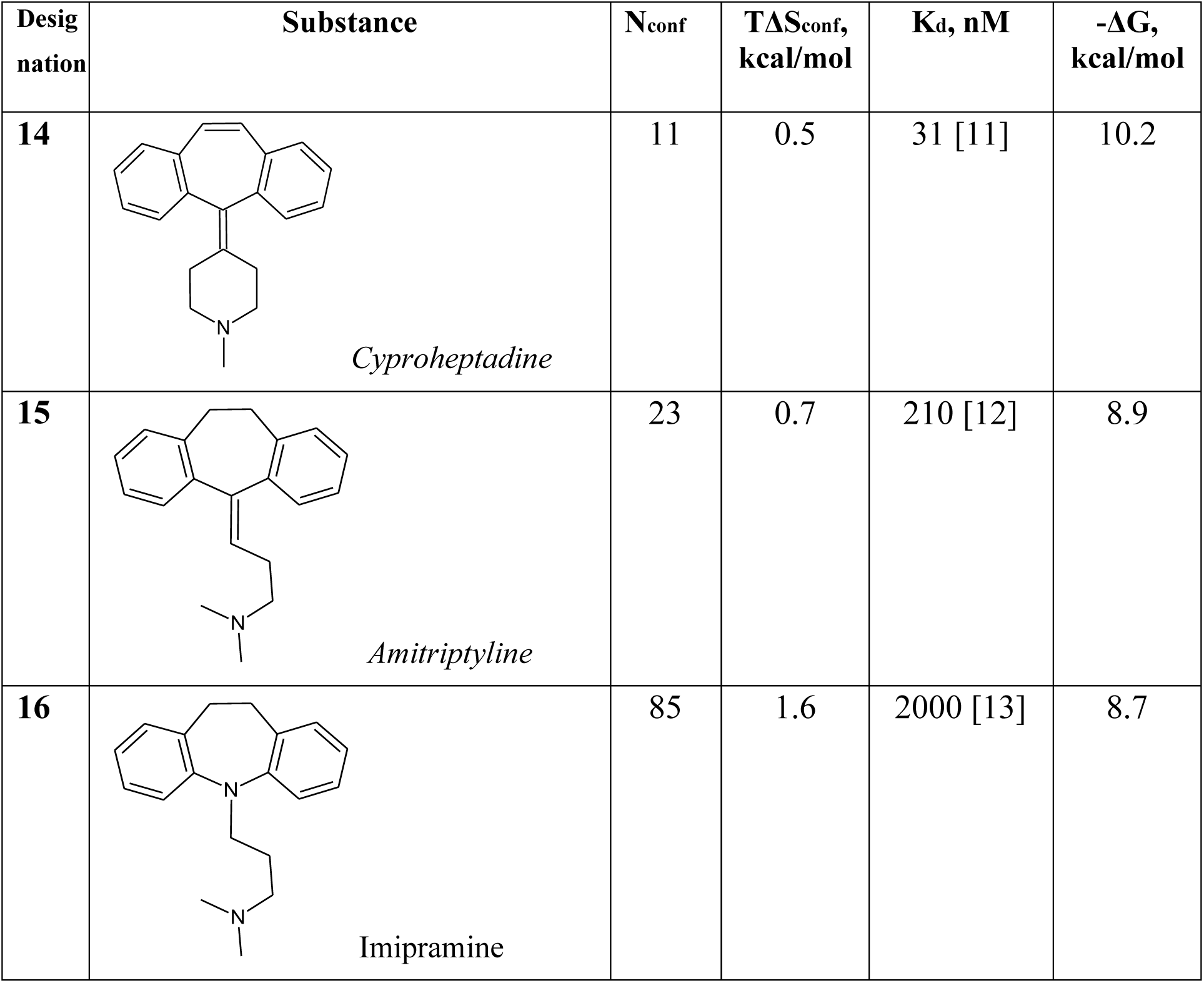
Conformational entropy of some antidepressants upon binding to D_2_-dopmaine receptors.

| Designation | Substance | $N_{\text{conf}}$ | $T\Delta S_{\text{conf}}$, kcal/mol | $K_d$, nM | $-\Delta G$, kcal/mol |
| --- | --- | --- | --- | --- | --- |
| 14 | *Cyproheptadine* | 11 | 0.5 | 31 [11] | 10.2 |
| 15 | *Amitriptyline* | 23 | 0.7 | 210 [12] | 8.9 |
| 16 | *Imipramine* | 85 | 1.6 | 2000 [13] | 8.7 |

Substitutions of single for double bonds and of a methylene chain for a conformationally stable N-methyl-4-piperidyl group make the substances structurally rigid (Table. 2). These structural changes certainly induce decreasing of N_conf_ and TΔS_conf_, and the matching increase of substances’ affinity to D_2_-receptors.

### Conformational entropy and ligands of the μ-opioid receptor

Conformational entropy can be considered as a universal factor in the preservation of receptor’s functional activity.

Values of the affinity and estimation of entropy losses during binding between morphine and met-enkephalin with μ-opioid receptors mediating their analgesic effect are summarized in Table 3. Met-enkephalin is pentapeptide (YGGFM), acting as an endogenous μ-receptor agonist.

**Table 3.** TΔS_conf_ and ΔG values for met-enkephalin and morphine binding to rat brain μ-opiate receptors.

| Ligand | MW | $N_{\text{conf}}$ | $T\Delta S_{\text{conf}}$,<br>kcal/mol | $K_d$, nM | $-\Delta G$,<br>kcal/mol |
| --- | --- | --- | --- | --- | --- |
| Met-enkephalin | 574 | 13329 | 3.84 | 3.0 [14] | 11.6 |
| Morphine | 286 | 5 | 0.57 | 1.5 [15] | 12.0 |

Meth-enkephalin with twice-higher molecular weight than morphine slightly differs in the affinity for μ-receptors. Like morphine and other opiates, met-enkephalin is competitively displaced from μ-receptors by _3_H-naloxone, and this is a proof for the common binding site for these substances at the receptor. Unlike rigid morphine molecule, met-enkephalin molecule is conformationally labile, having the increased N_conf_ and TΔS_conf_ values which produces affinity decrease. Previously we pointed that substances with K_d_ < 0.1 nM (ΔG < −14 kcal/mol) irreversibly inactivate receptor, so they become poisonous [2, 10]. It follows from Table 3 that enkephalins would be alike without conformational entropy influence.

Endogenous ligands of opioid receptors are not limited to enkephalins. It is endorphins which have more sizeable molecules. Thus, α- and β-endorphins with lengths of 16 and 31 amino acids are competitively displaced from μ-receptors by low-molecular-weight opiates and have relatively comparable affinities for these receptors 11 nM [13] and 3.2 nM [15], respectively.

Even if not all conformations are frozen in large molecules at their binding, conformational entropy plays the important and likely major role in preservation of receptor’s functional activity.

Little affinities of relatively large molecules are reasonably explained by steric hindrances to binding with receptor, and this is undoubtedly may be true. However, the found relation between Δ(TΔS_conf_) and ΔΔG indicates that entropy-based control of affinity is a major factor in the protection of neuroreceptors from irreversible inactivation at binding with large ligand molecules, such as enkephalins, endorphins, hormones, peptide drugs etc.

## Acknowledgments

We would like to make a statement of author contributions here. M.B.D. designed and performed calculations on conformational analysis and subsequent entropy changes, analyzed data and prepared the manuscript; F.S.D. analyzed data and wrote the paper.

